# General factors of white matter microstructure from DTI and NODDI in the developing brain

**DOI:** 10.1101/2021.11.29.470344

**Authors:** Kadi Vaher, Paola Galdi, Manuel Blesa Cabez, Gemma Sullivan, David Q Stoye, Alan J Quigley, Michael J Thrippleton, Debby Bogaert, Mark E Bastin, Simon R Cox, James P Boardman

## Abstract

Preterm birth is closely associated with diffuse white matter dysmaturation inferred from diffusion MRI and neurocognitive impairment in childhood. Diffusion tensor imaging (DTI) and neurite orientation dispersion and density imaging (NODDI) are distinct dMRI modalities, yet metrics derived from these two methods share variance across tracts. This raises the hypothesis that dimensionality reduction approaches may provide efficient whole-brain estimates of white matter microstructure that capture (dys)maturational processes. To investigate the optimal model for accurate classification of generalised white matter dysmaturation in preterm infants we assessed variation in DTI and NODDI metrics across 16 major white matter tracts using principal component analysis and structural equation modelling, in 79 term and 141 preterm infants at term equivalent age. We used logistic regression models to evaluate performances of single-metric and multimodality general factor frameworks for efficient classification of preterm infants based on variation in white matter microstructure. Single-metric general factors from DTI and NODDI capture substantial shared variance (41.8-72.5%) across 16 white matter tracts, and two multimodality factors captured 93.9% of variance shared between DTI and NODDI metrics themselves. General factors associate with preterm birth and a single model that includes all seven DTI and NODDI metrics provides the most accurate prediction of microstructural variations associated with preterm birth. This suggests that despite global covariance of dMRI metrics in neonates, each metric represents information about specific (and additive) aspects of the underlying microstructure that differ in preterm compared to term subjects.

**Highlights:**

- We measured variation of 7 DTI and NODDI metrics across 16 major tracts
- General factors for DTI and NODDI capture substantial shared variance across tracts
- General factors also capture substantial shared variance between DTI and NODDI
- Single-metric and multimodality factors associate with gestational age at birth
- The best preterm prediction model contains all 7 single-metric g-factors

## 1 Introduction

Diffusion tensor imaging (DTI) and neurite orientation dispersion and density imaging (NODDI) enable inference about the microstructural properties (such as water content, axonal density and myelination) of developing white matter from diffusion magnetic resonance imaging (dMRI) (Counsell et al., 2019; Tariq et al., 2016; Zhang et al., 2012). Neonatal dMRI has been valuable in assessing the impact of preterm birth on the developing brain; it reveals a preterm brain phenotype at term-equivalent age, which includes lower fractional anisotropy (FA) and neurite density index (NDI) and increased mean diffusivity (MD) throughout the white matter compared to term-born controls, with a dose-dependent effect of prematurity (Alexandrou et al., 2014; Anjari et al., 2007; Barnett et al., 2018; Batalle et al., 2017; Blesa et al., 2020; Boardman and Counsell, 2020; Hüppi et al., 1998; Partridge et al., 2004; Pogribna et al., 2013). Importantly, the dysconnectivity and reduced white matter integrity associated with preterm birth is substantially a whole brain phenomenon (Girault et al., 2019; Telford et al., 2017). This motivates the search for efficient whole-brain estimates that would capture maturational processes in early life.

Studies have demonstrated that dMRI measures of white matter tracts across the brain are correlated (e.g. an individual with high FA in one tract is likely to have high FA across other tracts) and that this relationship is exists across the life course (Cox et al., 2016; Girault et al., 2019; Lee et al., 2017; Mishra et al., 2013; Telford et al., 2017; Wahl et al., 2010). This property has allowed the derivation of general factors (g-factors) of white matter microstructure (e.g. gFA), which associate with general cognitive functioning (Alloza et al., 2016; Cox et al., 2019; Penke et al., 2010) and age (Cox et al., 2016). Similar diffusion properties have been observed in early life and these predict cognitive abilities (Lee et al., 2017). Our group has previously reported that in neonates DTI-metric-based g-factors explain around 50% of variance in eight white matter tracts and associate with gestational age (GA) at birth (Telford et al., 2017).

The different dMRI metrics themselves as well as the derived g-factors are correlated (Chamberland et al., 2019; Cox et al., 2016; De Santis et al., 2014; Girault et al., 2019; Penke et al., 2010), suggesting that dMRI measures share overlapping information which can cause partial redundancies in data analysis. Recently, a dimensionality reduction framework based on multimodal principal component analysis (PCA) was proposed (Chamberland et al., 2019; Geeraert et al., 2020). Using this framework the authors identified a small number of microstructurally informative and biologically-interpretable components/factors which captured 80% of variance in dMRI and myelin-sensitive imaging metrics across the white matter tracts, which associated with age in a sample of typically developing 8-18-year-old children (Chamberland et al., 2019; Geeraert et al., 2020).

In this work, using a neonatal dataset and a neonatal white matter tract atlas based on established protocols (Pecheva et al., 2017; Wakana et al., 2007), we aimed to: (1) determine g-factors for DTI and NODDI metrics and evaluate whether a single factor captures substantial variance across major tracts; and (2) investigate the shared variance across dMRI metrics by deriving a multimodal g-factor from DTI and NODDI, and quantify its predictive utility for GA at birth beyond uni-modal models. We hypothesised that g-factors would associate with GA at birth and that they would provide an efficient method for classifying generalised variation in white matter microstructure associated with preterm birth.

## 2 Materials and methods

### 2.1 Participants

Preterm (with GA at birth *<* 33 weeks) and term born infants were recruited as part of a longitudinal study (Theirworld Edinburgh Birth Cohort, TEBC) designed to investigate the effects of preterm birth on brain structure and long term outcome (Boardman et al., 2020). Exclusion criteria were major congenital malformations, chromosomal abnormalities, congenital infection, overt parenchymal lesions (cystic periventricular leukomalacia, haemorrhagic parenchymal infarction) or post-haemorrhagic ventricular dilatation. Ethical approval has been obtained from the National Research Ethics Service, South East Scotland Research Ethics Committee (11/55/0061, 13/SS/0143 and 16/SS/0154). Informed consent was obtained from a person with parental responsibility for each participant. The study was conducted according to the principles of the Declaration of Helsinki.

### 2.2 Data acquisition

Infants were scanned at the Edinburgh Imaging Facility: Royal Infirmary of Edinburgh, University of Edinburgh, UK using a Siemens MAGNETOM Prisma 3 T MRI clinical scanner (Siemens Healthcare Erlangen, Germany). A 16-channel phased-array paediatric head coil was used to acquire 3D T2-weighted SPACE images (T2w) (voxel size = 1mm isotropic, TE = 409 ms and TR = 3200 ms) and axial dMRI data. Diffusion MRI images were acquired in two separate acquisitions to reduce the time needed to re-acquire any data lost to motion artifacts: the first acquisition consisted of 8 baseline volumes (b = 0 s/mm^2^ [b0]) and 64 volumes with b = 750 s/mm^2^; the second consisted of 8 b0, 3 volumes with b = 200 s/mm^2^, 6 volumes with b = 500 s/mm^2^ and 64 volumes with b = 2500 s/mm^2^. An optimal angular coverage for the sampling scheme was applied (Caruyer et al., 2013). In addition, an acquisition of 3 b0 volumes with an inverse phase encoding direction was performed. All dMRI images were acquired using single-shot spin-echo echo planar imaging (EPI) with 2-fold simultaneous multislice and 2-fold in-plane parallel imaging acceleration and 2 mm isotropic voxels; all three diffusion acquisitions had the same parameters (TR/TE 3400/78.0 ms). Images affected by motion artifacts were re-acquired as required; dMRI acquisitions were repeated if signal loss was seen in 3 or more volumes. Infants were fed and wrapped and allowed to sleep naturally in the scanner. Pulse oximetry, electrocardiography and temperature were monitored. Flexible earplugs and neonatal earmuffs (MiniMuffs, Natus) were used for acoustic protection. All scans were supervised by a doctor or nurse trained in neonatal resuscitation.

### 2.3 Data preprocessing

Diffusion MRI processing was performed as follows: for each subject the two dMRI acquisitions were first concatenated and then denoised using a Marchenko-Pastur-PCA-based algorithm (Veraart et al., 2016); the eddy current, head movement and EPI geometric distortions were corrected using outlier replacement and slice-to-volume registration (Andersson et al., 2017, 2016, 2003; Andersson and Sotiropoulos, 2016); bias field inhomogeneity correction was performed by calculating the bias field of the mean b0 volume and applying the correction to all the volumes (Tustison et al., 2010). The T2w images were processed using the minimal processing pipeline of the developing human connectome project (dHCP) to obtain the bias field corrected T2w and the brain mask (Makropoulos et al., 2018, 2014). Finally, the mean b0 EPI volume of each subject was co-registered to their structural T2w volume using boundary-based registration (Greve and Fischl, 2009).

NODDI and DTI maps were calculated in the dMRI processed images to obtain: fractional anisotropy (FA), mean, axial and radial diffusivities (MD, AD and RD), neurite density index (NDI), isotropic volume fraction (ISO) and orientation dispersion index (ODI). To calculate the tensor derived metrics, only the first shell was used. NODDI metrics were calculated using the recommended values for neonatal white matter of the parallel intrinsic diffusivity (1.45 µm^2^/ms) (Guerrero et al., 2019; Zhang et al., 2012).

### 2.4 Tract segmentation

Whole brain tractography was performed in the ENA50 neonatal template space (Blesa et al., 2020, 2016) using SingleTensorFT tool within DTI-TK (Zhang et al., 2007, 2006) which generated white matter tractography from the ENA50 atlas tensor volume. Segmentation of white matter tracts was performed within the ENA50 atlas. Regions of interest (ROIs) used to delineate the tracts were drawn manually on the FA image, using the protocols outlined in Wakana et al. (2007) and Pecheva et al. (2017). Placement of ROIs is described in Supplementary Table 1 and these were drawn using the Paintbrush mode in ITK-SNAP (Yushkevich et al., 2006) (http://www.itksnap.org/). The ROIs were used to filter whole brain tractography either to select or to exclude tracts crossing the ROIs using TractTool within DTI-TK. The resulting tract images were binarized and manually refined. The white matter tracts delineated are shown in Figure 1.

**Figure 1:**
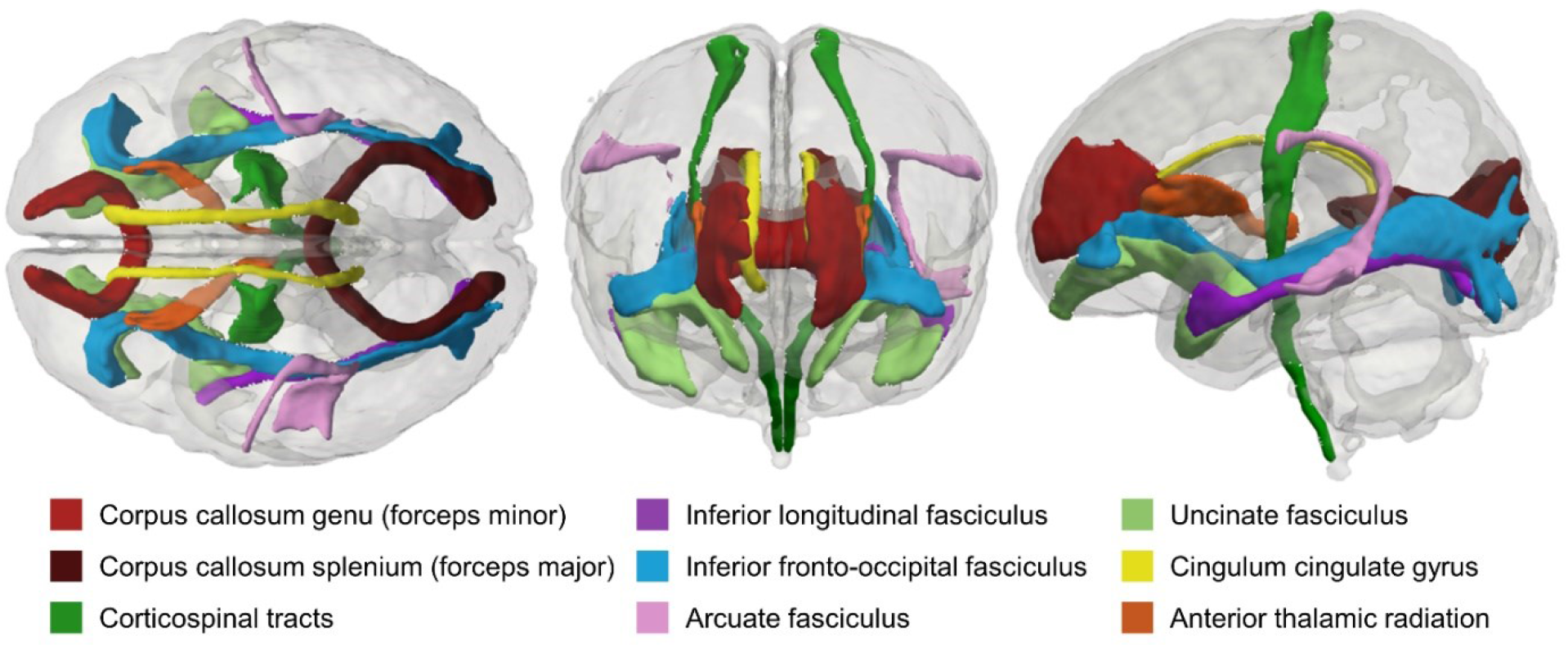
Visual representation of the generated white matter tracts in the ENA50 neonatal atlas space. Shown in superior (left), anterior (centre) and lateral (right) views.

### 2.5 Tract segmentation in subjects’ native space and extraction of tract-averaged dMRI metrics

T2w processed images were registered to the ENA50 T2w structural template using rigid, affine and symmetric normalization (SyN) (Avants et al., 2008). The resulting transformation was concatenated with the previously computed transformation from B0 to T2w and used to bring the tract ROIs defined in the ENA50 space to each subject’s native space in a single step.

The average multi-tissue response function was calculated across the full population (Dhollander et al., 2019, 2016; Smith et al., 2020), with a FA threshold of 0.1. Then, the multi-tissue fibre orientation distribution (FOD) was calculated (Jeurissen et al., 2014) with the average response function using a spherical harmonic order (L_max_) of 8. Only two (white matter and cerebrospinal fluid) response functions were used. Finally, a joint bias field correction and multi-tissue informed log-domain intensity normalisation on the FODs images was performed (Raffelt et al., 2017).

The tracts in native space were created using the iFOD2 algorithm (Tournier et al., 2010). The propagated tract ROIs were dilated and the original tract ROIs were used as seed images for the tractography, while the dilated tract ROIs were used as masks to constrain the tracts. The length of the fibres was set with a minimum length of 20 mm and a maximum of 250 mm. Finally, for each tract, a track density image (TDI) map (number of tracts per voxel) was created and normalized between 0 and 1 (Calamante et al., 2010). For each tract, the TDI map was multiplied by each of the DTI and NODDI maps, summed and divided by the average of the TDI map to calculate the weighted tract-averages for each of the DTI and NODDI metrics.

### 2.6 Statistical analysis

All statistical analyses were performed in R (version 4.0.5) (R Core Team, 2020).

#### 2.6.1 Effect of preterm birth on tract-averaged dMRI metrics

The tract-averaged dMRI parameters were adjusted for GA at scan by fitting a linear model of each scaled (z-transformed) metric on GA at scan and retaining the residuals. The distributions of the residualised dMRI metrics in each tract were assessed for normality using the Shapiro–Wilk test. Student’s t-test or Mann– Whitney U-test as a non-parametric alternative was used to compare the tract-averaged values between term and preterm groups; Spearman’s rho was used to investigate correlations between tract-averaged values and GA at birth. Reported p-values were adjusted for the false discovery rate (FDR) using the Benjamini-Hochberg procedure (Benjamini and Hochberg, 1995).

#### 2.6.2 Single-metric g-factors

The average Pearson’s correlation coefficient for the inter- and intra-hemispheric associations between the tracts was calculated by first transforming the Pearson’s r values to Fisher’s Z, taking the average, and then back-transforming the value to Pearson’s correlation coefficient (Corey et al., 1998).

One PCA was conducted for each of the seven DTI (FA, MD, AD, RD) and NODDI (NDI, ODI, ISO) parameters across the 16 tracts to quantify the proportion of shared variance between them. Thus, in each analysis, each subject was described by 16 features, computed as the tract-averaged values of a given metric across each tract. The first unrotated principal component (PC) scores were extracted as the single-metric g-factors. The g-factors were adjusted for GA at scan by fitting a linear model of each g-factor on GA at scan and retaining the residuals. We report regression coefficients for linear models fitting a linear each of the residualised g-factors and GA at scan. All values were scaled (z-transformed) before fitting the models, thus, the regression coefficients are in the units of standard deviations. Reported p-values were adjusted for the FDR using the Benjamini-Hochberg procedure.

Structural equation modelling was used to investigate the extent that differences in GA at birth explain the shared variance across tracts (a common pathway model where GA has associations with only the latent g-factor), and the extent that GA at birth conveys unique information about individual tracts that is not conveyed via shared variance. First, we evaluated the similarities between the g-factors obtained using PCA with the measurement model (confirmatory factor analysis [CFA] within the structural equation model) that was conducted for each metric using the R package *lavaan* (Rosseel, 2012). We used full information maximum likelihood estimation. Model fit was assessed according to standard fit indices: χ^2^ test, root mean square error of approximation (RMSEA), comparative fit index (CFI), Tucker-Lewis index (TLI), and standardised root mean square residual (SRMR). Residual covariance paths (paths linking specific tracts to one another to account for the specific similarities between related tracts beyond their shared covariance across all tracts) were added between each of the bilateral tracts, the genu and splenium of the corpus callosum, as well as anatomically overlapping tracts (Dice Coefficient > 0.1 based on the dilated tract masks in template: ILF and IFOF in the same hemisphere, IFOF and UNC in the same hemisphere, ILF and UNC in the same hemisphere, and splenium of the corpus callosum and bilateral IFOF). Pearson’s correlation coefficients were calculated for the g-factors derived using PCA and CFA.

Thereafter, we tested three models where 1) GA has associations with only the latent g-factor – a common pathway model; 2) GA has associations with each of the individual tracts separately and not with the latent factor – an independent pathways model; and 3) GA is associated with the latent factor and also with some specific factors – a common + independent pathways model (Cox et al., 2016; Tucker-Drob, 2013). To estimate the common + independent pathways model, we first included a path from GA to the latent g-factor, and then, in an iterative fashion, used modification indices (with a minimum value of 10) to include any additional paths from GA to specific tracts that substantially improved model fit. All models were adjusted for GA at scan at g-factor level. See Supplementary Figure 1 for graphical representation of the structural equation models. We used the *χ*^2^ difference test (*aov* function within *lavaan*) and model fit indices (Akaike Information Criterion [AIC], Bayesian Information Criterion [BIC], and sample size adjusted Bayesian Information Criterion [saBIC]) to examine the fit differences between the models.

#### 2.6.3 Multimodal g-factor

A multimodal PCA was conducted by pooling all tracts and metrics using a modification of an established framework (Chamberland et al., 2019; Geeraert et al., 2020). In summary, all metrics were analysed together in a single PCA, so that each observation was an individual tract described by the 7 dMRI metrics, for a total of *n*×*t* observations, where *n* is the number of subjects and *t* is the number of tracts. The first and second PC were extracted as the multimodal g-factors which were averaged across the 16 tracts for each participant. To study the effect of GA at birth on the multimodal g-factors, we first adjusted the g-factors for GA at scan by fitting a linear model of each g-factor on GA at scan and retaining the residuals; then, linear regression models were fitted for each of the residualised multimodal g-factors and GA at birth. All values were scaled (z-transformed) before fitting the models, thus, the regression coefficients are in the units of standard deviations. Reported p-values were adjusted for the FDR using the Benjamini-Hochberg procedure.

#### 2.6.4 Prediction modelling

We used the single- and multimodal g-factors as predictors in logistic regression models to discriminate between preterm and full-term infants. We measured classification accuracy using a 10-repeated 10-fold cross-validation scheme. In each of 10 repetitions data were randomly split in 10-folds of which 9-folds were used as training set to compute the PCs, adjust these for GA at scan, and train the prediction of preterm vs term subjects. The g-factors in the test set were computed and adjusted for GA at scan using the models retained from the training set. Then, the generalisation ability of the logistic regression model to predict term vs preterm group trained on the training set was assessed in the test set. Folds were stratified to preserve the proportion of term and preterm subjects of the whole sample. Accuracy was computed as the percentage of correctly classified subjects across folds and repetitions. We estimated the empirical distribution of chance by repeating the prediction analysis 1000 times after randomly assigning each subject to either the preterm or term group; permutation p-values were calculated by counting how many times the null models obtained an accuracy equal or greater than the original model.

### 2.7 Data and code availability

Reasonable requests for original image and anonymised data will be considered through the BRAINS governance process (www.brainsimagebank.ac.uk) (Job et al., 2017). The segmented tracts in the ENA50 template space are available here: https://git.ecdf.ed.ac.uk/jbrl/ena. The code for tract propagation and average calculation, as well as scripts for the data analysis in this paper are available here: https://git.ecdf.ed.ac.uk/jbrl/neonatal-gfactors.

## 3 Results

### 3.1 Study sample

The study group consisted of 220 neonates: 141 participants were preterm and 79 were term-born controls. Demographic details for participant characteristics are provided in Table 1. Among the preterm infants, 30 (21.3%) had bronchopulmonary dysplasia (defined as need for supplementary oxygen ≥36 weeks GA), 7 (5%) developed necrotising enterocolitis requiring medical or surgical treatment, and 27 (19.1%) had an episode of postnatal sepsis defined as either blood culture positivity with a pathogenic organism, or physician decision to treat for ≥5 days in the context of growth of coagulase negative staphylococcus from blood or a negative culture.

**Table 1:** Neonatal participant characteristics. The last column reports the p-values of the group differences computed with t-test for continuous variables and Fisher’s exact test for categorical variables.

|  | term (n=79) | preterm (n=141) | term vs. preterm |
| --- | --- | --- | --- |
| GA at birth (weeks) | 39.65 (36.42 - 42.14) | 29.48 (23.42 - 32.94) | n/a |
| Birth weight (grams) | 3482 (2410 - 4560) | 1334 (500 - 2510) | n/a |
| Birth weight z-score | 0.48 (-2.30 - 2.57) | -0.02 (-3.13 - 2.14) | $p < 0.001$ |
| GA at scan (weeks) | 42.07 (38.28 - 46.14) | 40.78 (36.56 - 45.84) | $p < 0.001$ |
| M:F ratio | 43:36 | 83:58 | $p = 0.571$ |

GA = Gestational age, M = male, F = female.

### 3.2 Associations between preterm birth and tract-averaged dMRI metrics

Figure 2 and Supplementary Table 2 show tract-averaged dMRI parameter values for each of the 16 tracts for the term and preterm neonates. After adjusting for GA at MRI, in the majority of tracts FA was lower and MD, RD, AD and ISO were higher in preterm infants compared to term-born controls. However, ATR, CCG and CST showed only minimal or no differences in the DTI metrics between the two groups. There were groupwise differences in tract-averaged NDI and ODI values in a minority of the tracts (Figure 2).

**Figure 2:**
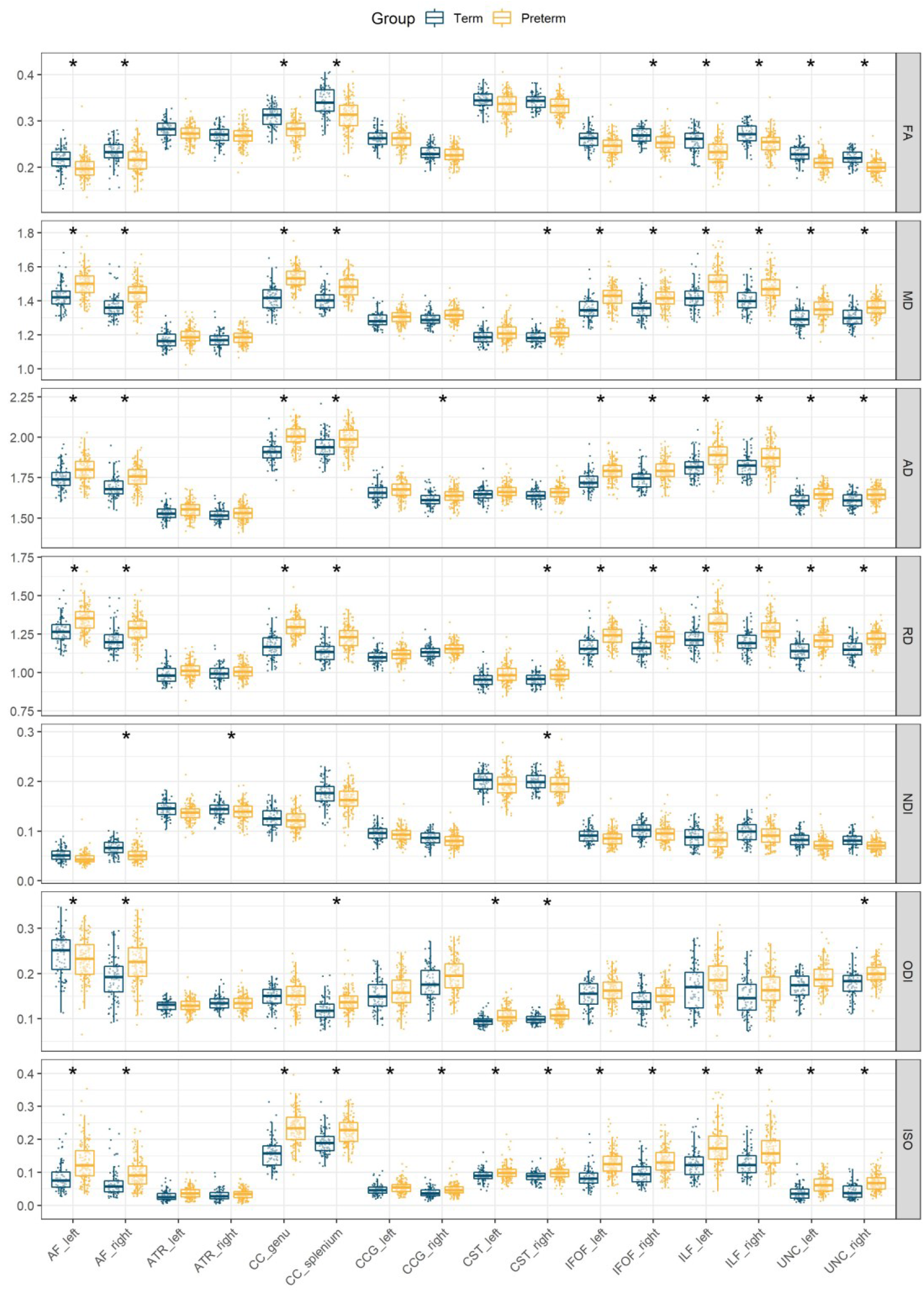
Tract-averaged diffusion characteristics of brain white matter tracts. Asterisks (*) indicate statistically significant (FDR-corrected p*<*0.05) differences in tract-averaged values between term and preterm infants after adjusting for age at scan. FA = fractional anisotropy, MD = mean diffusivity, AD = axial diffusivity, RD = radial diffusivity, NDI = neurite density index, ODI = orientation dispersion index, ISO = isotropic volume fraction, CC genu = corpus callosum genu/forceps minor, CC splenium = corpus callosum splenium/forceps major, CST = corticospinal tract, IFOF = inferior fronto-occipital fasciculus, ILF = inferior longitudinal fasciculus, AF = arcuate fasciculus, UNC = uncinate fasciculus, CCG = cingulum cingulate gyrus, ATR = anterior thalamic radiation

### 3.3 Single-metric general factors of white matter microstructure

For all DTI and NODDI metrics, with the exception of ODI, metrics across tracts correlate positively (Figure 3). The mean (±SD) of the correlations was 0.601 (±0.294) for FA, 0.713 (±0.217) for MD, 0.573 (±0.199) for AD, 0.721 (±0.234) for RD, 0.628 (±0.250) for NDI, 0.351 (±0.287) for ODI, and 0.584 (±0.221) for ISO.

**Figure 3:**
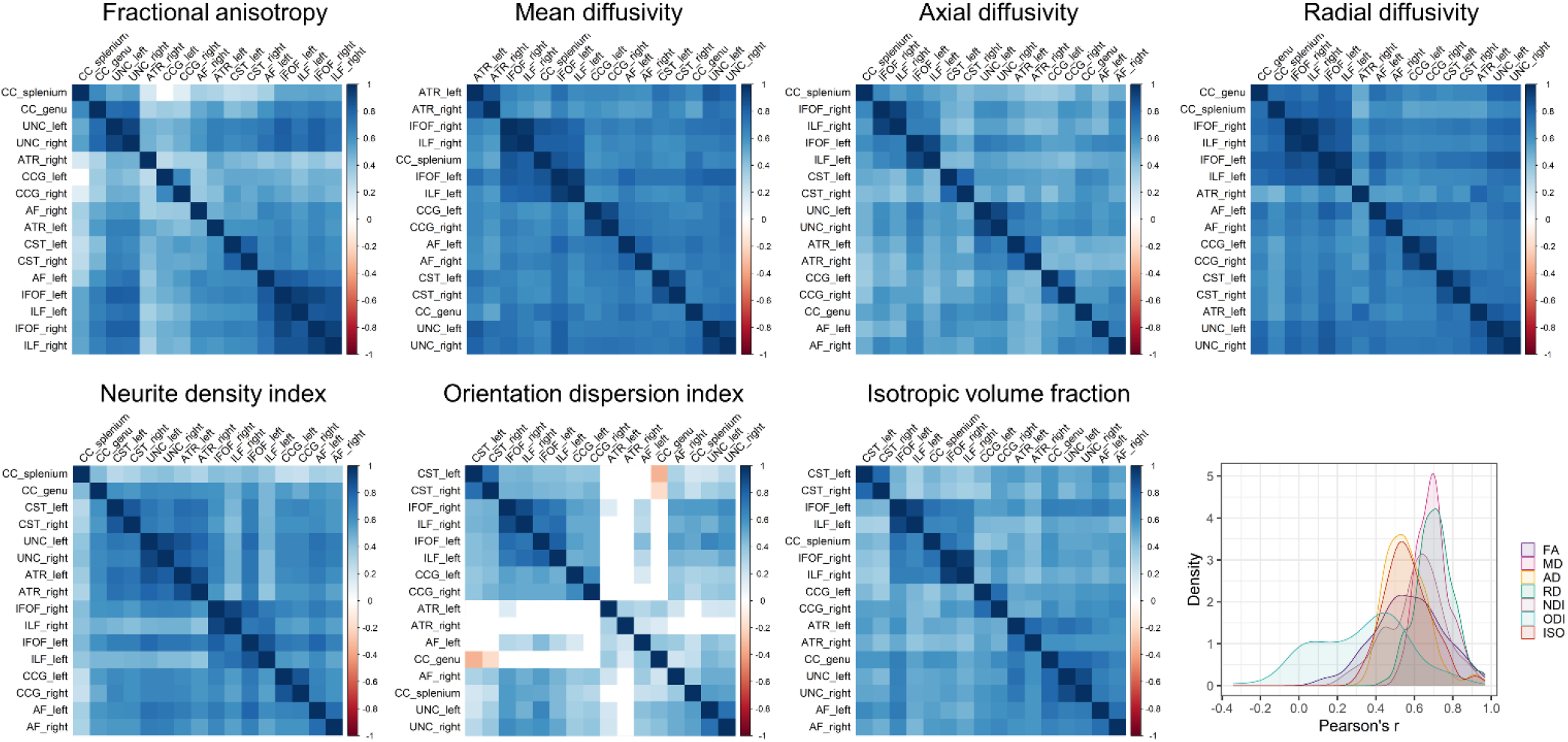
Heatmaps of inter- and intra-hemispheric associations (Pearson’s r) for tract-averaged DTI (top row) and NODDI (bottom row) metrics. In each case, the heatmaps are arranged by grouping highly correlated tracts around the diagonal. Blank squares represent correlations that were not nominally statistically significant (p*>*0.05). The plot on the bottom right represents the density of the correlation magnitudes. CC genu = corpus callosum genu/forceps minor, CC splenium = corpus callosum splenium/forceps major, CST = corticospinal tract, IFOF = inferior fronto-occipital fasciculus, ILF = inferior longitudinal fasciculus, AF = arcuate fasciculus, UNC = uncinate fasciculus, CCG = cingulum cingulate gyrus, ATR = anterior thalamic radiation.

We conducted separate PCAs for each of the 7 DTI and NODDI metrics on 16 white matter tracts to derive single-metric g-factors. For each metric, the scree plot provided evidence for a strong single factor capturing common variance across the tracts indicated by the comparatively large eigenvalue (Fig. 4). This was less clear for ODI, which had a weaker first component and stronger second component compared to other dMRI metrics. The first PC is the g-factor for each of the white matter diffusion measures and this explained 61.3% variance in FA, 71.9% in MD, 59.9% in AD, 72.6% in RD, 63.9% in NDI, 41.8% in ODI, and 59.8% in ISO across the tracts. The tract loadings for the single-metric g-factors are presented in Table 2.

**Figure 4:**
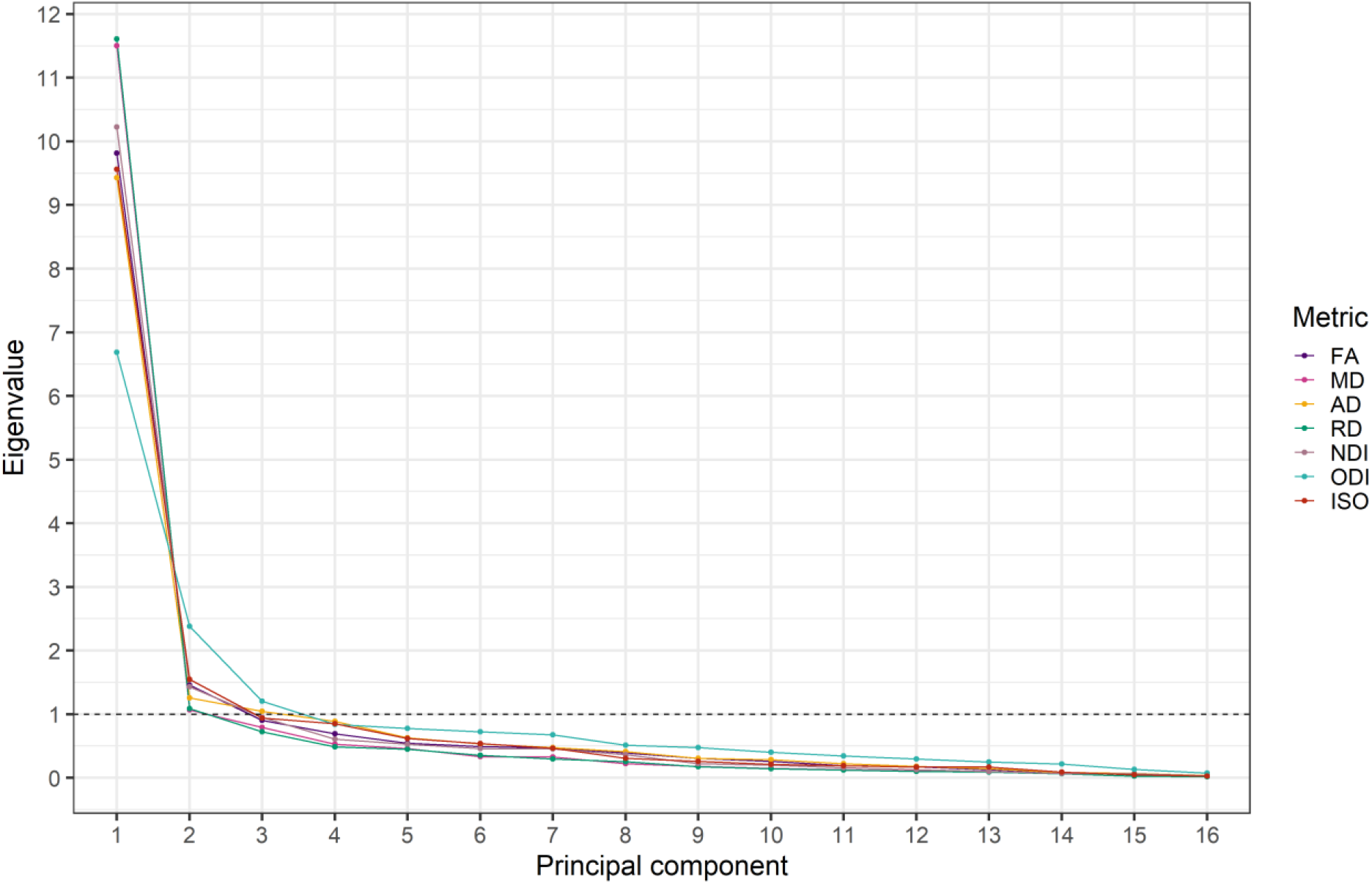
Scree plot for the principal component analysis, showing the eigenvalue against the number of components for each white matter tract dMRI metric. FA = fractional anisotropy, MD = mean diffusivity, AD = axial diffusivity, RD = radial diffusivity, NDI = neurite density index, ODI = orientation dispersion index, ISO = isotropic volume fraction.

**Table 2:**
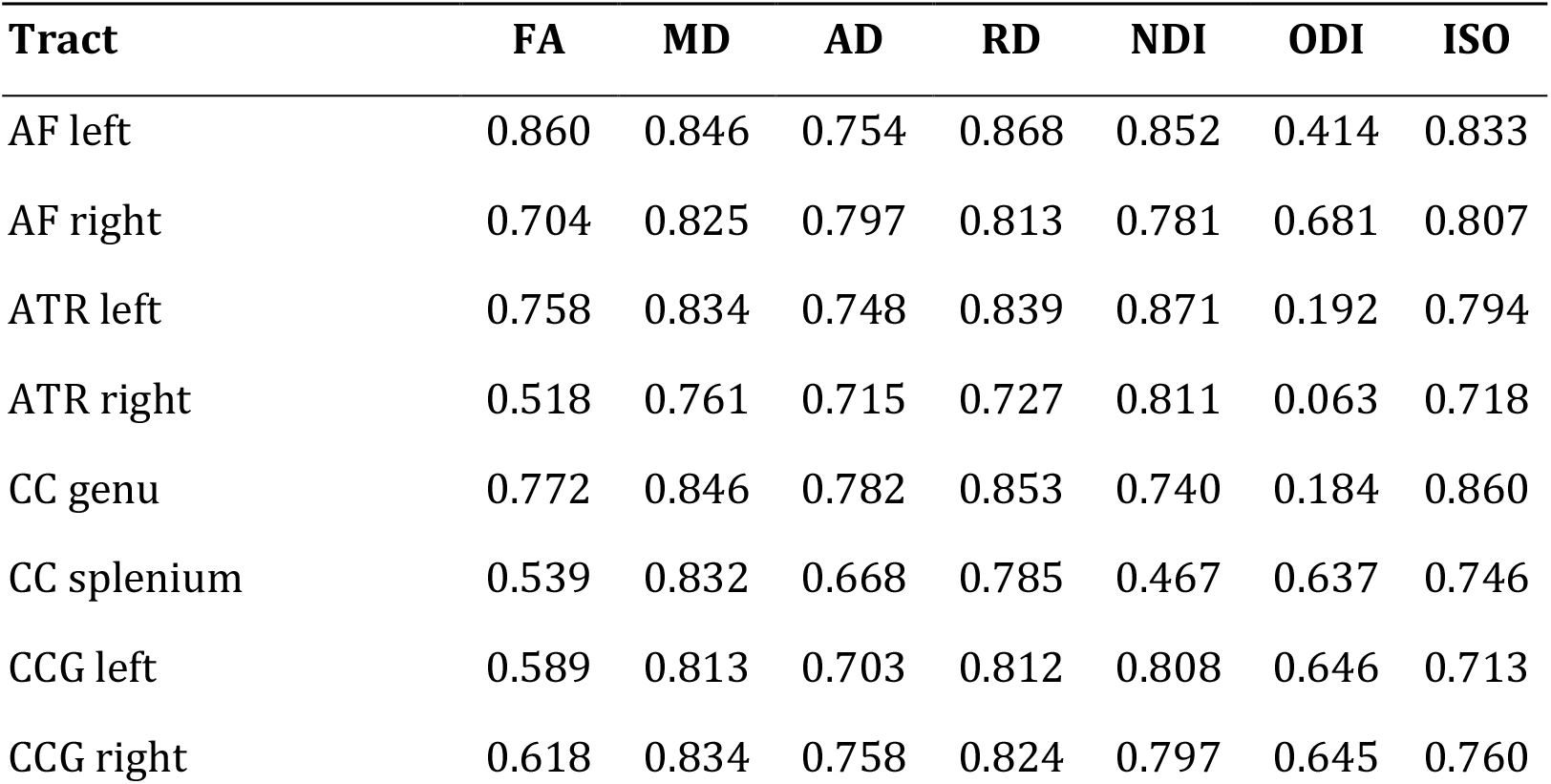
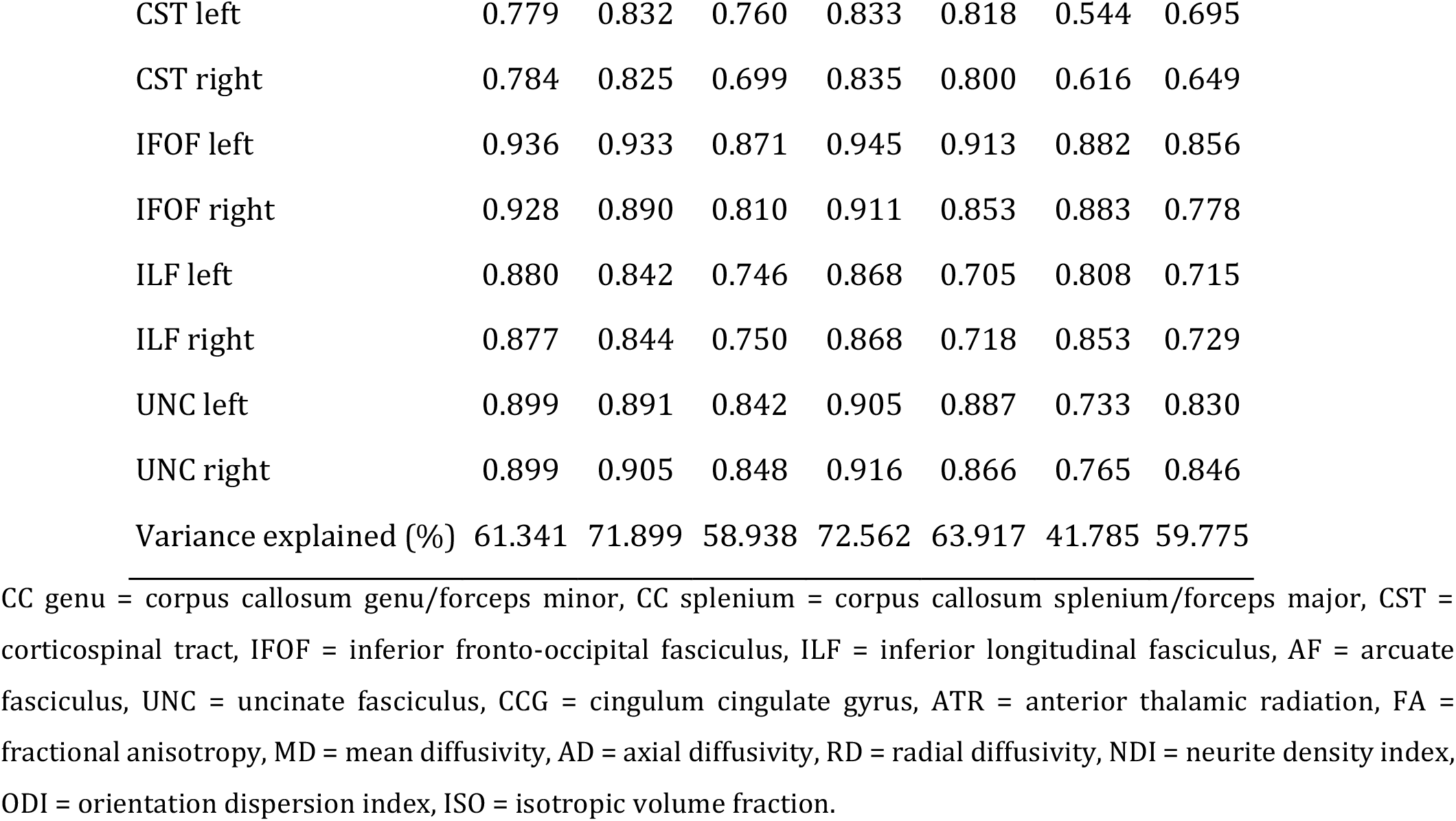
Tract loadings (correlation between the manifest variable and extracted component score) and explained variance for the first unrotated principal component (g-factor) for the seven dMRI metrics.

| Tract | FA | MD | AD | RD | NDI | ODI | ISO |
| --- | --- | --- | --- | --- | --- | --- | --- |
| AF left | 0.860 | 0.846 | 0.754 | 0.868 | 0.852 | 0.414 | 0.833 |
| AF right | 0.704 | 0.825 | 0.797 | 0.813 | 0.781 | 0.681 | 0.807 |
| ATR left | 0.758 | 0.834 | 0.748 | 0.839 | 0.871 | 0.192 | 0.794 |
| ATR right | 0.518 | 0.761 | 0.715 | 0.727 | 0.811 | 0.063 | 0.718 |
| CC genu | 0.772 | 0.846 | 0.782 | 0.853 | 0.740 | 0.184 | 0.860 |
| CC splenium | 0.539 | 0.832 | 0.668 | 0.785 | 0.467 | 0.637 | 0.746 |
| CCG left | 0.589 | 0.813 | 0.703 | 0.812 | 0.808 | 0.646 | 0.713 |
| CCG right | 0.618 | 0.834 | 0.758 | 0.824 | 0.797 | 0.645 | 0.760 |
| CST left | 0.779 | 0.832 | 0.760 | 0.833 | 0.818 | 0.544 | 0.695 |
| CST right | 0.784 | 0.825 | 0.699 | 0.835 | 0.800 | 0.616 | 0.649 |
| IFOF left | 0.936 | 0.933 | 0.871 | 0.945 | 0.913 | 0.882 | 0.856 |
| IFOF right | 0.928 | 0.890 | 0.810 | 0.911 | 0.853 | 0.883 | 0.778 |
| ILF left | 0.880 | 0.842 | 0.746 | 0.868 | 0.705 | 0.808 | 0.715 |
| ILF right | 0.877 | 0.844 | 0.750 | 0.868 | 0.718 | 0.853 | 0.729 |
| UNC left | 0.899 | 0.891 | 0.842 | 0.905 | 0.887 | 0.733 | 0.830 |
| UNC right | 0.899 | 0.905 | 0.848 | 0.916 | 0.866 | 0.765 | 0.846 |
| Variance explained (%) | 61.341 | 71.899 | 58.938 | 72.562 | 63.917 | 41.785 | 59.775 |

After adjustment for GA at scan, there were significant associations between GA at birth and general factors of FA, MD, AD, RD and ISO (Figure 5). The strongest relationship was seen between GA at birth and gISO (GA at birth explained 11.06% of variance in gISO). Interestingly, GA at birth did not significantly associate with the g-factors of biophysical measures of white matter microstructure (NDI and ODI), mirroring the single-tract results described above.

**Figure 5:**
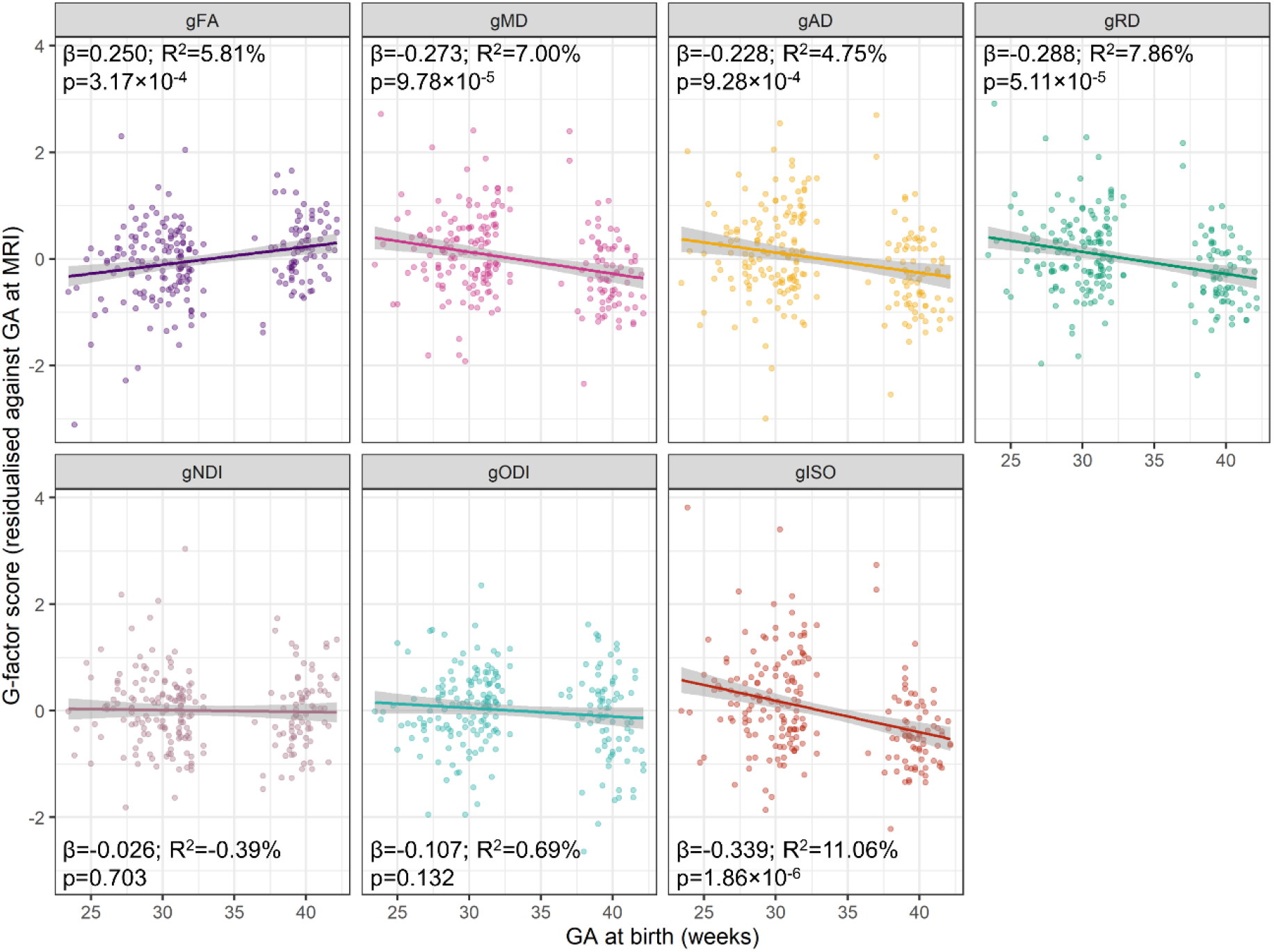
Associations between GA at birth and the g-factors of the seven dMRI metrics. Regression lines and 95% confidence intervals (shaded) are shown for linear regression models between GA at birth and the g-factor scores, adjusted for GA at scan. The *β* coefficients are in standardised units so represent a standard deviation change in the residualised g-factor scores per standard deviation increase in GA at birth; variance explained in the model is shown in adjusted R^2^. Reported p-values are adjusted for the false discovery rate (FDR) using the Benjamini-Hochberg procedure. FA = fractional anisotropy, MD = mean diffusivity, AD = axial diffusivity, RD = radial diffusivity, NDI = neurite density index, ODI = orientation dispersion index, ISO = isotropic volume fraction.

To investigate the extent to which shared variance across all tracts explains differences in GA at birth, or whether specific tracts carry further information beyond generalised covariance, we used structural equation modelling. We observed that the measurement model (CFA) for each DTI and NODDI metric is highly collinear with the PCA results indicated by the similarities between the factor loadings (Supplementary Table 3; see Supplementary Table 4 for fit indices; all CFI > 0.93 (except ODI, CFI = 0.891)) and high positive correlations between the g-factors derived using PCA and CFA (all r>0.98; Supplementary Table 5).

The structural equation modelling results showed that for the general factors of FA, MD, AD, RD and ISO there was evidence that GA at birth significantly associated with the g-factor (common model). The independent pathway model (where GA at birth associates with the tract-specific values) fit significantly better than the model that only included the common pathway of GA at birth associations (Table 3; for factor loadings and regression coefficients see Supplementary Table 6), although it included the highest number of paths. We inspected the modification indices of the common pathway model to determine whether there are incremental, unique tract-specific effects of GA at birth which are not conveyed by the effect of GA at birth on the shared variance. The modification indices did not indicate additional tract-specific paths associated with GA at birth for any metric, thus, we were unable to construct models with both common and independent pathways and this suggests that the common model provides sufficient refinement of the properties of white matter microstructure that are affected by GA at birth.

**Table 3:**
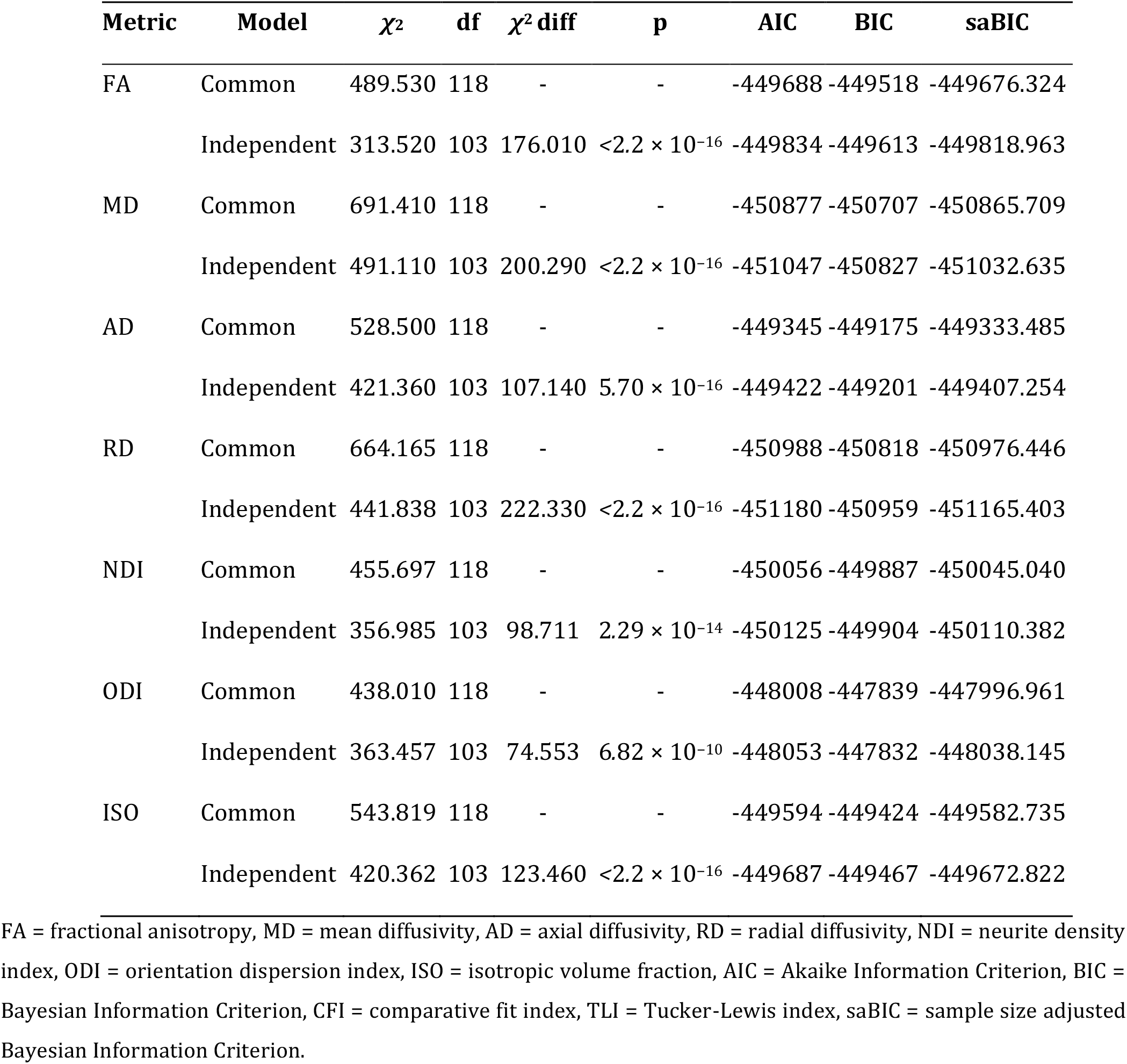
Model fit indices for each of the structural equation models linking GA at birth with the g-factors or individual white matter tracts. P-values refer to the difference (*χ*^2^ difference test) between the common and the independent pathway models. For full parameter estimates in these models see Supplementary Table 6.

### 3.4 Multimodal general factors of white matter microstructure

Next, we studied the shared variance of DTI and NODDI metrics across white matter tracts. The correlation matrices in Figure 5 show that the metrics form two clusters of positively correlated metrics: the first cluster represents positive correlations between FA and NDI, and the second cluster of positive correlations is formed of MD, RD, AD and ISO, while ODI appears to be a weaker member of the second cluster. These two clusters are negatively correlated with each other. However, there is also variability in between-metric correlations between the different tracts. Nevertheless, the correlation matrix in the middle panel of Figure 6 highlights the similarity between the microstructural measures, which is consistent with them representing shared information about tract microstructure.

**Figure 6:**
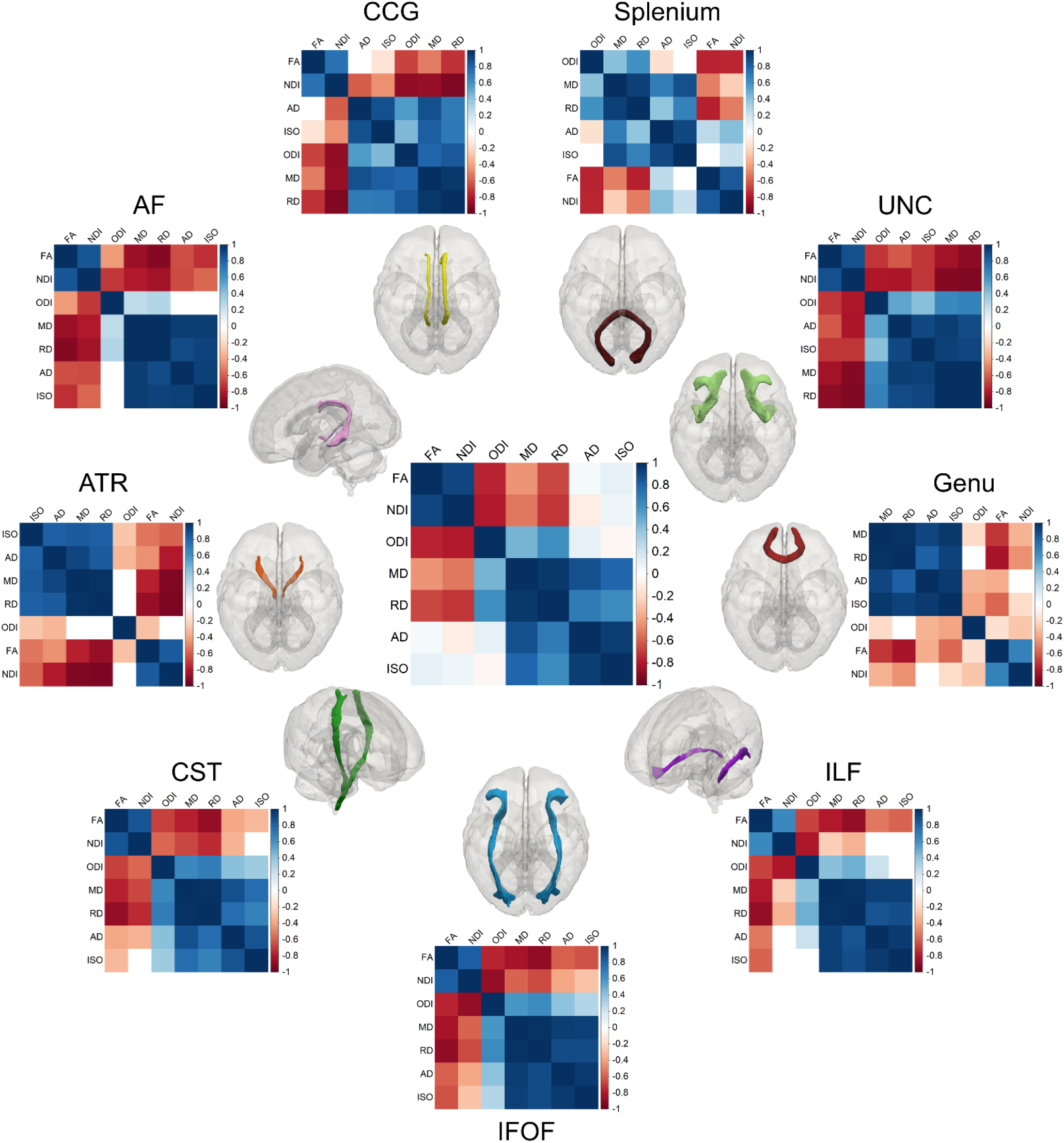
Correlation matrices of the seven diffusion measures. The middle image represents the average of all white matter tracts. Matrices are re-organised using hierarchical clustering, grouping measures that have similar correlations together. Note that for bilateral tracts, the left and right values were averaged prior to performing the correlation. Genu = corpus callosum genu/forceps minor, splenium = corpus callosum splenium/forceps major, CST = corticospinal tract, IFOF = inferior fronto-occipital fasciculus, ILF = inferior longitudinal fasciculus, AF = arcuate fasciculus, UNC = uncinate fasciculus, CCG = cingulum cingulate gyrus, ATR = anterior thalamic radiation, FA = fractional anisotropy, MD = mean diffusivity, AD = axial diffusivity, RD = radial diffusivity, NDI = neurite density index, ODI = orientation dispersion index, ISO = isotropic volume fraction.

A PCA including all seven DTI and NODDI metrics revealed that 93.9% of the variability in dMRI metrics across white matter tracts is accounted by the first two PCs (Figure 6). The first PC (proportion of variance explained 60.0%, *λ* = 4.20) is mostly composed of RD and MD (both contributing negatively, 23.4% and 21.4%, respectively), and the second PC which captures 33.9% of variance in the data (*λ* = 2.38) is mostly driven by ISO (26.8%), AD (21.3%) and FA (18.9%). The loadings and contributions of the dMRI metrics to the first two PCs are presented in Table 4. RD and MD appear to be solely loading onto the PC1 (together contribute *<*5% to the PC2), while the other dMRI metrics have more similar contributions to PC1 and PC2. The variability of between-tract correlations of dMRI metrics as mentioned above is also reflected in the clustering of tracts on the PC axes (Figure 7).

**Figure 6:**
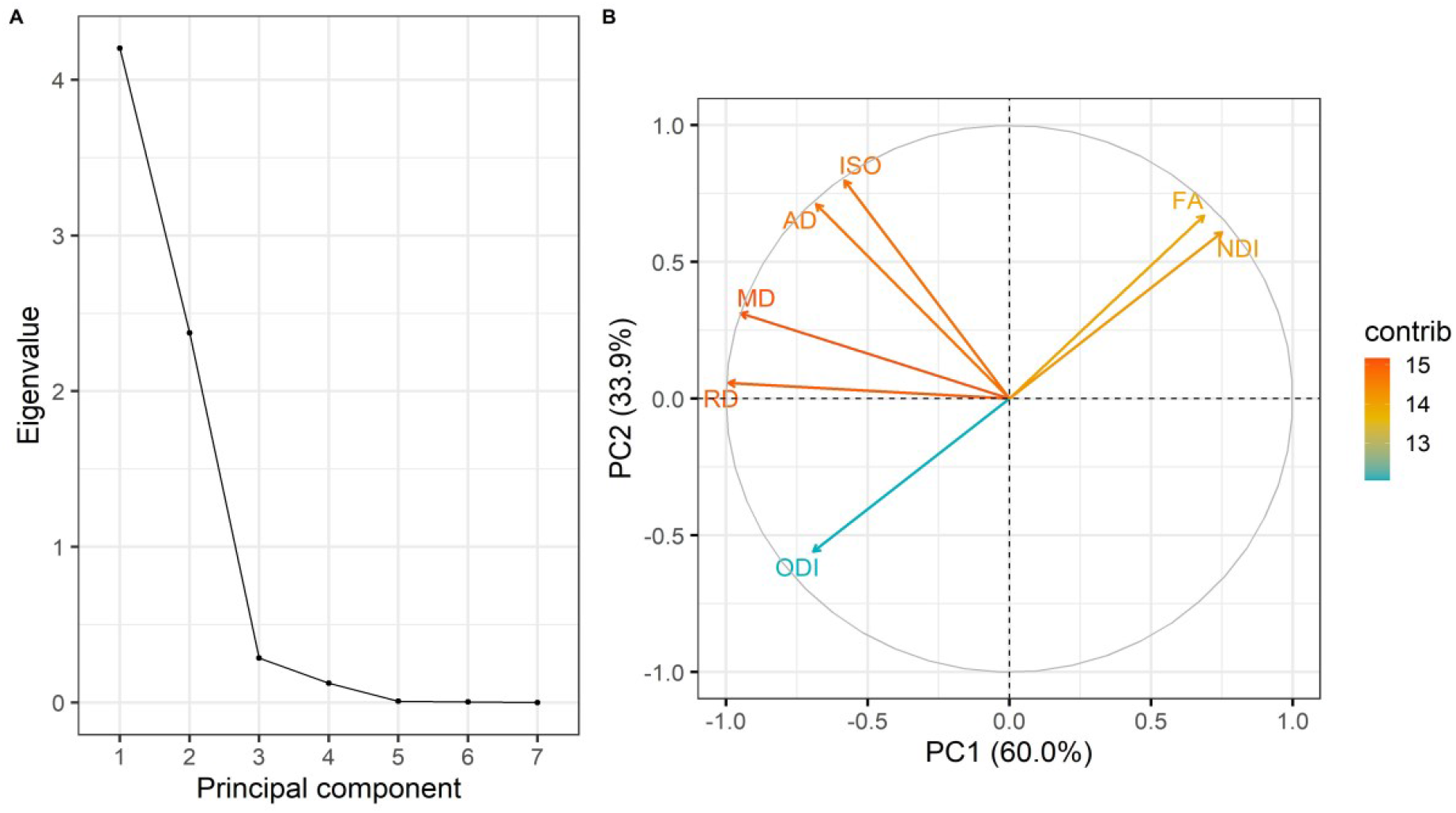
Multimodal PCA. (A) Scree plot of the eigenvalues, (B) PCA variable contribution plot; the colours represent the contribution of the dMRI metric to the components. FA = fractional anisotropy, MD = mean diffusivity, AD = axial diffusivity, RD = radial diffusivity, NDI = neurite density index, ODI = orientation dispersion index, ISO = isotropic volume fraction.

**Table 4:**
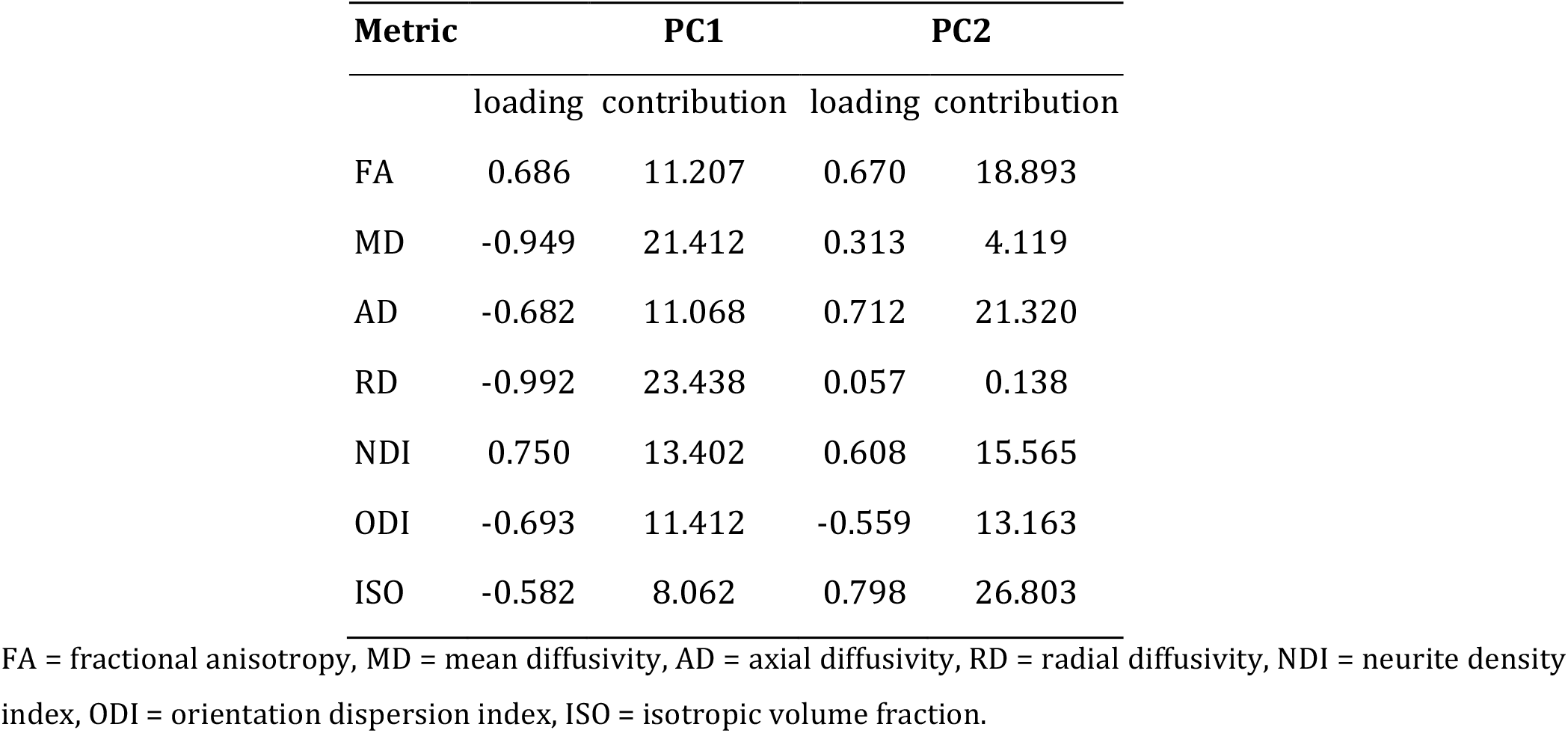
dMRI metric loadings to the multimodal principal components.

**Figure 7:**
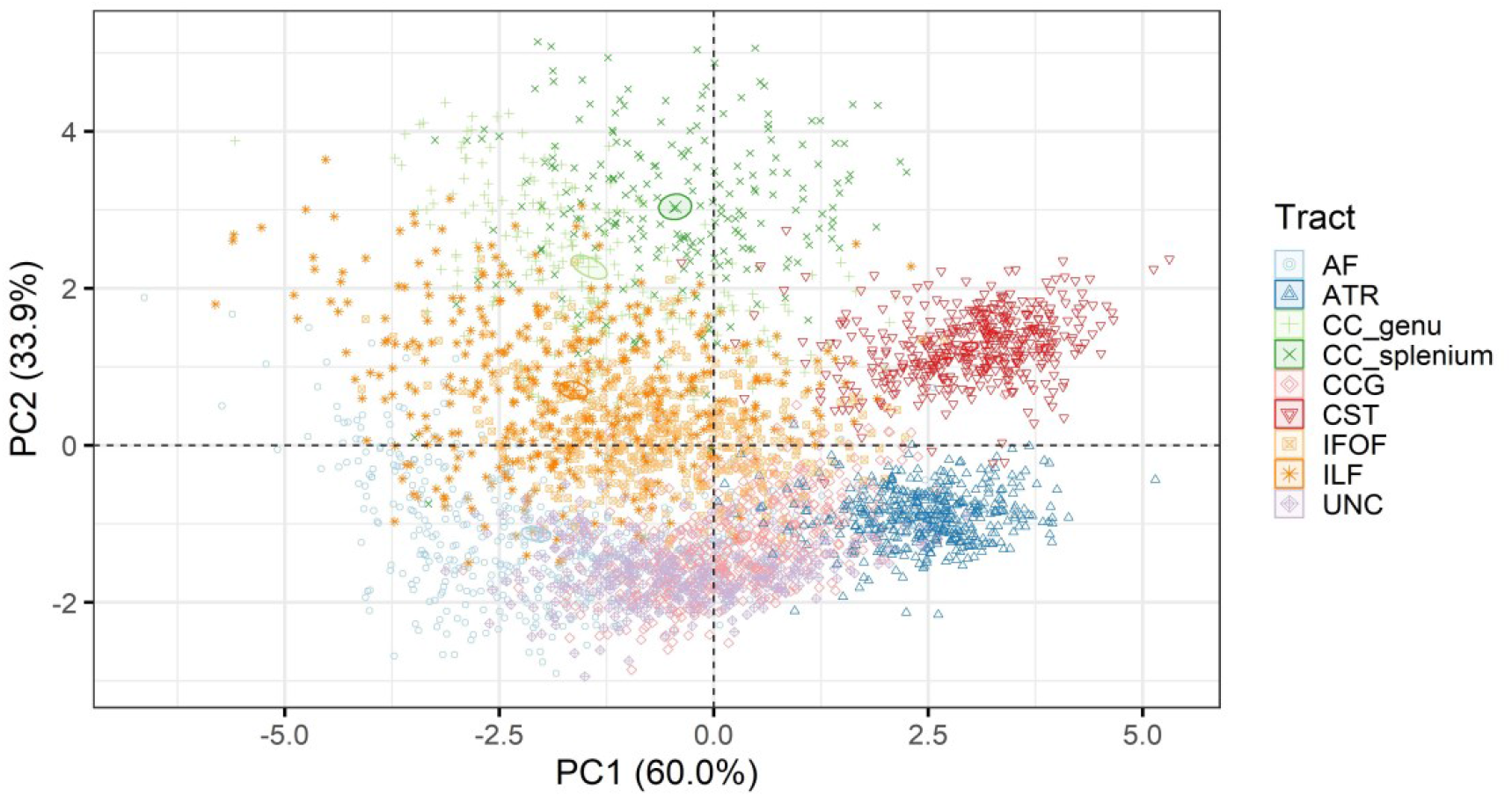
Visualisation of individual tract coordinates on the multimodal principal component axes. CC genu = corpus callosum genu/forceps minor, CC splenium = corpus callosum splenium/forceps major, CST = corticospinal tract, IFOF = inferior fronto-occipital fasciculus, ILF = inferior longitudinal fasciculus, AF = arcuate fasciculus, UNC = uncinate fasciculus, CCG = cingulum cingulate gyrus, ATR = anterior thalamic radiation

GA at birth was significantly associated with both multimodal g-factors (Figure 8): the first multimodal g-factor that has high negative contributions from RD and MD was positively, and the second multimodal g-factor with high positive contributions from ISO and AD was negatively associated with GA at birth. It is possible that individual tracts may contribute to varying degrees to the relationship with age (Supplementary Table 2).

**Figure 8:**
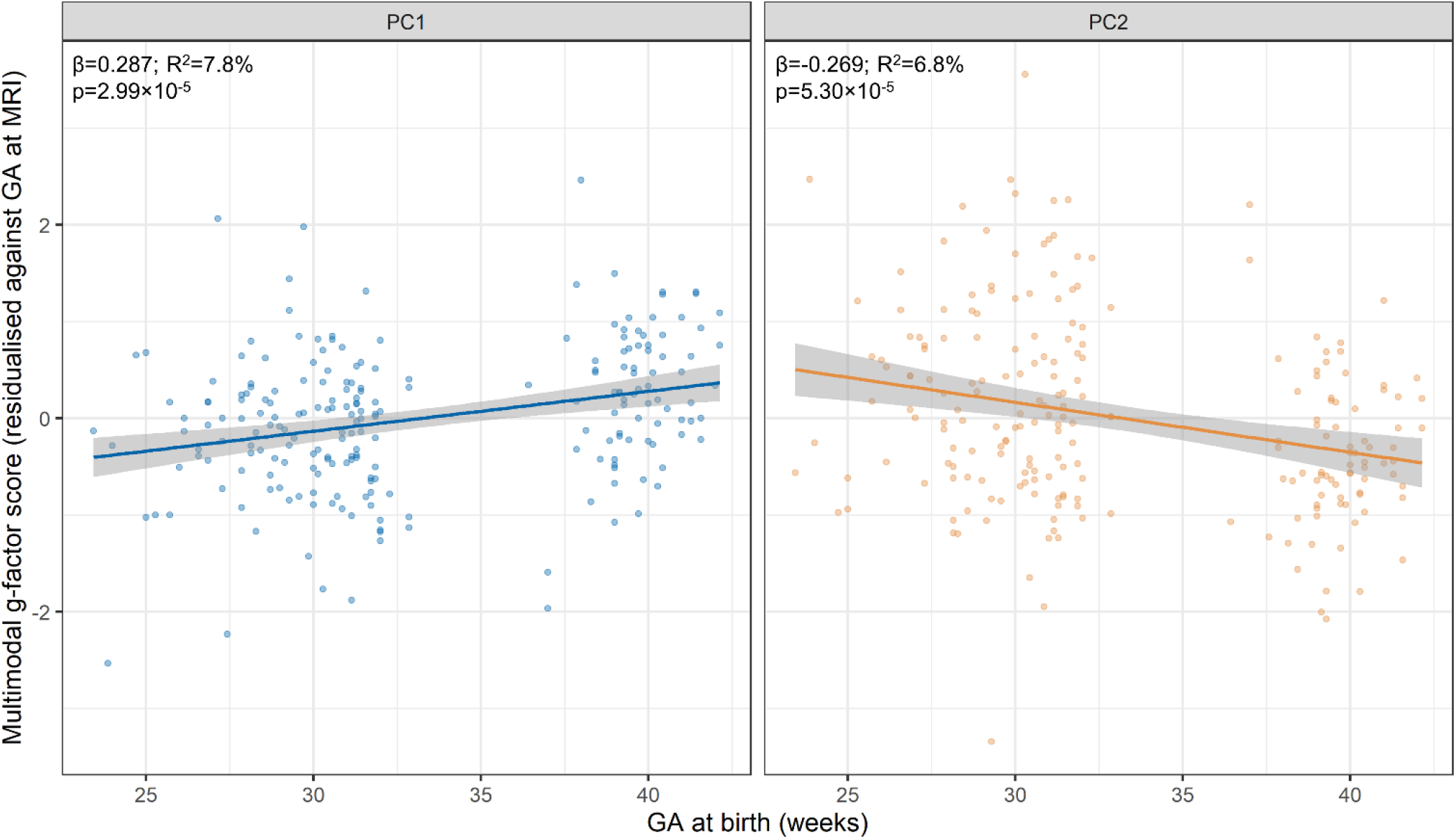
Associations between GA at birth and the multimodal g-factors. The extracted multimodal principal components were averaged across the 16 tracts for each participant which resulted in a single estimate for the multimodal g-factors for each subject. Regression lines and 95% confidence intervals (shaded) are shown for linear regression models between GA at birth and the g-factor scores, adjusted for GA at scan. The *β* coefficients are in standardised units so represent a standard deviation change in the residualised g-factor scores per standard deviation increase in GA at birth; variance explained in the model is shown in adjusted R^2^. Reported p-values are adjusted for the false discovery rate (FDR) using the Benjamini-Hochberg procedure.

### 3.5 Utility of g-factors to classify infants based on gestational age

Given the high shared variance within and between the dMRI metrics across major white matter tracts and the significant associations between GA at birth and the derived g-factors, we asked whether g-factors are able to classify infants based on GA at birth (preterm vs term classification) (Table 5). Overall, the prediction accuracy for the single metric and multimodal g-factors only marginally exceeded chance (64.1%). The highest prediction accuracy (75.2%) was achieved when incorporating all single metric g-factors in one model, however, it has to be noted that this is the least parsimonious model with seven predictors compared to one and two in the other models.

**Table 7:** Prediction model results based on 10-repeated 10-fold cross validated logistic regression models using the single-metric g-factors and multimodal g-factors. Reported values are mean and standard deviations computed across cross-validation folds and repetitions. Permutation p-values are computed over 1000 random permutations of the group variable.

|  | Accuracy | Permutation p-value |
| --- | --- | --- |
| gFA | 64.9±5.0 | 0.005 |
| gMD | 67.3±6.4 | 0 |
| gRD | 67.6±7.1 | 0 |
| gAD | 65.5±6.6 | 0 |
| gNDI | 64.1±1.0 | 0.210 |
| gODI | 64.0±3.2 | 0.703 |
| gISO | 67.7±8.0 | 0 |
| All DTI | 65.5±7.6 | 0.002 |
| All NODDI | 71.3±8.3 | 0 |
| All single-metric | 75.2±8.4 | 0 |
| Multimodal PC1 | 67.9±6.9 | 0 |
| Multimodal PC2 | 64.3±7.8 | 0.010 |
| Multimodal PC1 and PC2 | 67.5±8.5 | 0 |

FA = fractional anisotropy, MD = mean diffusivity, AD = axial diffusivity, RD = radial diffusivity, NDI = neurite density index, ODI = orientation dispersion index, ISO = isotropic volume fraction, DTI = diffusion tensor imaging, NODDI= neurite orientation dispersion and density imaging.

## 4 Discussion

In this study, utilising the substantial shared variance within and between DTI and NODDI metrics across 16 major white matter tracts, we derive single- and multimetric g-factors which covary with GA at birth. Using structural equation modelling, we show that whilst the shared variance among tracts carries much of the white matter microstructural information about GA-based differences, there is modest additional unique information at the level of individual pathways that enhances term/preterm differentiation, though larger samples are required to reliably estimate the precise magnitudes and loci of the most informative white matter pathways. We demonstrate that combining single-metric g-factors from DTI and NODDI together in one prediction model offered the most efficient method for characterising variation in white matter microstructure associated with preterm birth, suggesting each metric carries additive information. These results add to the body of literature suggesting generalised dysmaturation of the white matter in the preterm neonates.

Variance in measures derived from dMRI is shared among white matter tracts in both neonates and adults, and previous studies have suggested that generalised measures to capture global white matter microstructure can be derived (Cox et al., 2016; Lee et al., 2017; Penke et al., 2010; Telford et al., 2017). Here, we report that the g-factors capture 58.9-72.6% of variance in DTI metrics, thus replicating our previous results in an independent, larger sample of neonates and different tract segmentation protocol (Telford et al., 2017). We additionally expand on the previous work and report that similarly to DTI metrics, in neonates there is substantial shared variance of NODDI metrics across white matter tracts (41.8-62.9%) as was previously reported in adult population (Cox et al., 2016). We observed the largest variance captured by a single g-factor for RD, while there was least evidence for a single latent factor for ODI, indicated by the comparably smaller eigenvalue for the first component, suggesting that this measure of white matter microstructure may be capturing tract-specific rather than global effects.

The correlations between different DTI and NODDI metrics themselves indicate that they share overlapping information in the brain (Chamberland et al., 2019; De Santis et al., 2014), but less is known about the covariance of dMRI measures in early development when water diffusion properties are different. By examining the covariance of dMRI metrics averaged over 16 white matter tracts, we observed that there are two clusters of positively correlated metrics: the first cluster includes measures of microstructural complexity/integrity of FA and NDI while the second cluster includes measures related to water diffusivity (MD, RD, AD and ISO); the metrics in these two clusters are in turn negatively correlated with one another. The highest positive correlations are between the pairs of FA-NDI (microstructural complexity/integrity), RD-MD (hindrance and degree of diffusivity) and AD-ISO (free/diffuse water). Importantly, the dMRI metric covariance structures vary slightly between tracts, confirming the tract-specific variability highlighted by the CFA. For example, the splenium of the corpus callosum appears to have weaker between-metric correlations overall although high correlations between MD-RD, FA-NDI and AD-ISO are still present. Interestingly, ODI, on average, appears to have weaker correlations with other dMRI metrics, which may further suggest between-tract variability of this measure of fibre orientation. Indeed, in the uncinate, inferior fronto-occipital fasciculi, cingulum cingulate gyri and corticospinal tracts, ODI is a part of the second cluster of positively correlated metrics, while it has very low correlations with other metrics in the inferior longitudinal fasciculi and the anterior thalamic radiation and is negatively associated with all other dMRI metrics in the genu of the corpus callosum. AD and ISO correlations with NDI, FA and ODI appear considerably weaker on average, possibly also due to variations in the dMRI metric covariance structure between tracts. Together, our results suggest that similarly to what is observed in adults and children (Chamberland et al., 2019; Geeraert et al., 2020), the interdependence of dMRI measures is already present at birth.

We found that a considerable proportion of variance is shared across the dMRI metrics in neonates, which confirms previous observations in children and adolescents (Chamberland et al., 2019; Geeraert et al., 2020). We used this shared variance to derive multimodal g-factors of white matter microstructure using PCA as a data reduction technique. The two extracted multimodal g-factors together explained almost 94% of variance in the seven DTI and NODDI metrics across 16 white matter tracts. Due to the different MRI measures used in the current work compared to those profiled by Chamberland et al. (2019) and Geeraert et al. (2020), we are unable to make direct comparisons in the interpretation of the multimodal g-factors. However, similarly to these previous papers, we also observed that the multimodal PCs generally consist of dMRI metrics that share similarities in their sensitivity to different tissue properties. The multimodal PC1 that accounts for the largest proportion of variance in the data (60%) consists of measures sensitive to hindrance/restricted water diffusion and magnitude of diffusivity (RD and MD). The multimodal PC2 accounting for 34% of variance in the data consists of measures of free water (ISO) as well as axonal integrity (AD, FA).

We were then interested in testing whether the derived g-factors can be used to characterise atypical white matter development associated with low GA. After adjusting for age at scan, we report that gFA was positively and gMD, gAD and gRD negatively associated with GA at birth. Thus, we replicate previous results in a larger independent cohort and across more tracts (Telford et al., 2017). In addition, here we report significant negative association between GA at birth and gISO, which had the strongest correlation with GA at birth among the DTI and NODDI g-factors. Thus, those infants born preterm exhibit less coherent, but a greater magnitude of water diffusion across the major white matter tracts in the brain compared to term-born controls. Interestingly, despite the substantial variance reported in NDI and ODI across white matter tracts, gNDI and gODI are not significantly associated with GA at birth. This could indicate that these two metrics capture more specific aspects of tract composition, which may be less meaningful at a global level.

These results together suggest generally lower white matter integrity and higher water diffusivity in infants born preterm compared to term, and are in line with findings obtained using other analysis approaches such as tract-based spatial statistics (Barnett et al., 2018; Thompson et al., 2019) or tract-specific analyses (Pecheva et al., 2017). We used structural equation modelling to test whether the common variance shared among all tracts is sufficient to explain differences between infants born at varying GA. We found that the tract-specific (independent pathways) model is significantly better than the common model, suggesting there is incrementally valid, low level information for GA at birth contained in the unique tract-specific microstructural properties. However, it has to be noted that this model included the highest number of paths, and the residual variance that cannot be accounted for by the common factor constitutes both tract-specific aspects of microstructure and measurement error. We could not reliably detect any additional tract-specific pathways to be substantially more informative for GA at birth compared to the general/common factor, suggesting that the most parsimonious model, in which GA at birth affects the global/shared variance of the tracts, offers valuable distillation of the between-person differences in white matter microstructure that are pertinent for GA variability.

The prediction modelling results revealed that the single-metric g-factors (except for gNDI and gODI) achieved preterm vs term classification accuracy significantly higher than chance, but the classification accuracy was relatively low. It could be hypothesised that preterm birth has a diffuse effect on white matter microstructure, which is better captured by methods that do not rely on anatomically constrained regions (e.g. peak width of skeletonised metrics (Baykara et al., 2016; Blesa et al., 2020)). Nevertheless, the g-factors could carry information beyond the simple dichotomy of term vs preterm birth and could be useful for investigating other environmental or genetic/epigenetic exposures that are hypothesised to affect global white development (Boardman et al., 2014; Boardman and Counsell, 2020; Krishnan et al., 2017; Wheater et al., 2021), or for predicting neurocognitive outcomes as previously reported in adults and children (Cox et al., 2019, 2016; Lee et al., 2017; Penke et al., 2010).

We also report that the multimodal g-factors associate with GA at birth, which, given the correlations of the dMRI metrics with the multimodal g-factors, give a similar interpretation of the effect of GA at birth on dMRI metrics. However, despite the significant association, the preterm vs term classification accuracy achieved using the multimodal g-factors was, similarly to single-metric g-factors, relatively low. Interestingly, however, we achieved the greatest classification accuracy when combining all single metric g-factors together in one prediction model. These results may imply that despite global covariance of dMRI metrics in neonates, each one carries information on specific (and additive) aspects of the underlying microstructure that differ in preterm compared to term subjects. It is important to acknowledge that the model combining all single metric g-factors is by far the least parsimonious model tested, and increasing the number of predictors could artificially inflate the estimation of prediction accuracy. However, the combined single g-factor prediction model is by far the most successful one and we have used cross-validation with the aim to minimise bias and militate against the artificial inflation.

## 5 Conclusion

In this work, we extracted tract-averaged DTI and NODDI metrics from 16 major white matter tracts in 220 neonates of wide-ranging GA at birth. We then applied PCA as a data reduction technique to derive single- and multimodal general factors of white matter microstructure. These g-factors explained substantial variance within and between DTI and NODDI metrics across white matter tracts and associated with GA at birth. Combining single-metric g-factors together in one prediction model achieved discriminating power between term and preterm infants. This framework may be useful for investigating the upstream determinants and neurocognitive consequences of diseases characterised by atypical white matter development.

## 6 CRediT author statement

**Kadi Vaher**: Conceptualization, Methodology, Formal analysis, Investigation, Data Curation, Writing Original Draft, Visualization, Funding acquisition; **Paola Galdi**: Conceptualization, Methodology, Formal analysis, Writing - Original Draft; **Manuel Blesa Cabez**: Methodology, Software, Writing - Original Draft; **Gemma Sullivan**: Investigation, Resources, Writing - Review & Editing; **David Q Stoye**: Investigation, Resources, Writing - Review & Editing; **Alan J Quigley**: Investigation, Resources; **Michael J Thrippleton**: Methodology, Software, Resources, Writing - Review & Editing; **Debby Bogaert**: Conceptualization, Supervision, Writing - Review & Editing; **Mark E Bastin**: Methodology, Software, Validation, Writing - Review & Editing; **Simon R Cox**: Conceptualization, Methodology, Writing - Review & Editing; **James P Boardman**: Conceptualization, Methodology, Writing - Original Draft, Supervision, Funding acquisition.

## 8 Acknowledgments

We are grateful to the families who consented to take part in the study. Neonatal participants were scanned in the University of Edinburgh Imaging Research MRI Facility at the Royal Infirmary of Edinburgh which was established with funding from The Wellcome Trust, Dunhill Medical Trust, Edinburgh and Lothians Research Foundation, Theirworld, The Muir Maxwell Trust and many other sources. We are thankful to all the University’s imaging research staff for providing the infant scanning.

## Abbreviations

AD: axial diffusivity
AF: arcuate fasciculus
AIC: Akaike Information Criterion
ATR: anterior thalamic radiation
BIC: Bayesian Information Criterion
CC genu: corpus callosum genu/forceps minor
CC splenium: corpus callosum splenium/forceps major
CCG: cingulum cingulate gyrus
CFA: confirmatory factor analysis
CFI: comparative fit index
CST: corticospinal tract
dMRI: diffusion MRI
DTI: diffusion tensor imaging
FA: fractional anisotropy
FDR: false discovery rate
FOD: fibre orientation distribution
GA: gestational age
IFOF: inferior fronto-occipital fasciculus
ILF: inferior longitudinal fasciculus
ISO: isotropic volume fraction
MD: mean diffusivity
NDI: neurite density index
NODDI: neurite orientation dispersion and density imaging
ODI: orientation dispersion index
PCA: principal component analysis
RD: radial diffusivity
RMSEA: root mean square error of approximation
ROI: region of interest
SRMR: standardised root mean square residual
TDI: track density image
TLI: Tucker-Lewis index
UNC: uncinate fasciculus

