## Supplementary Figure 1 for "General factors of white matter microstructure from DTI and NODDI in the developing brain"

### Supplementary Figures

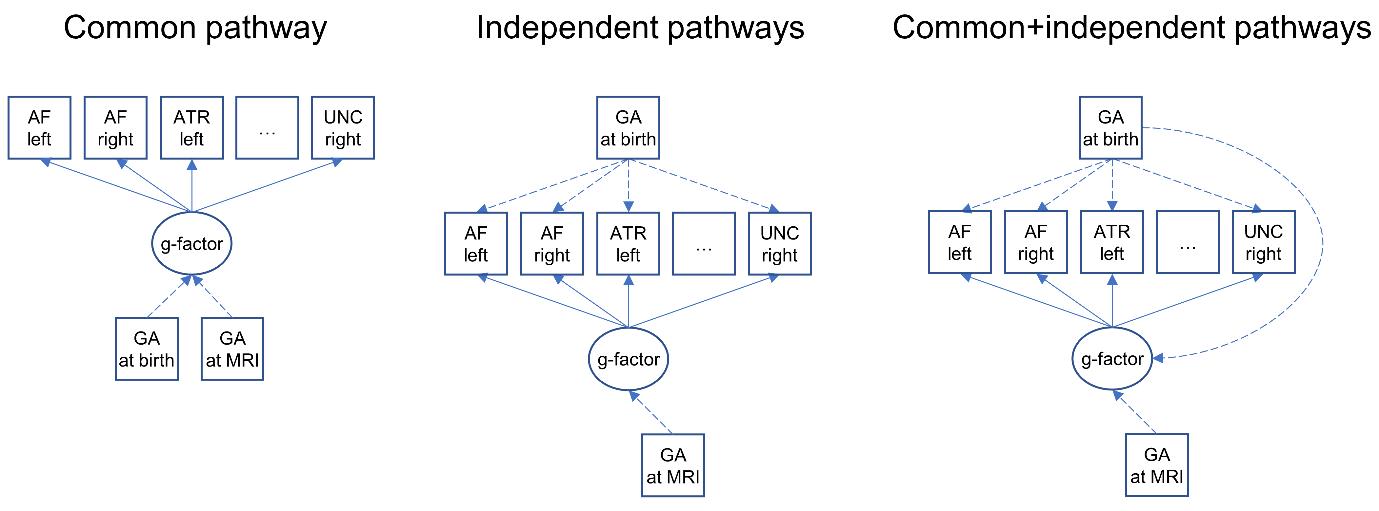

**Supplementary Figure 1.** Representation of the structural equation modelling approach. Factor loadings are represented with solid lines; dashed lines represent the regression coefficients. AF = arcuate fasciculus, UNC = uncinate fasciculus, ATR = anterior thalamic radiation, GA = gestational age.
